# Sea butterflies in a pickle: Reliable biomarkers and seasonal sensitivity of pteropods to ocean acidification in the Gulf of Maine

**DOI:** 10.1101/2023.07.31.551235

**Authors:** Amy E. Maas, Gareth L. Lawson, Alexander J. Bergan, Zhaohui Aleck Wang, Ann M. Tarrant

## Abstract

The passive dissolution of anthropogenically produced CO_2_ into the ocean system is reducing ocean pH and changing a suite of chemical equilibria, with negative consequences for some marine organisms, in particular those that bear calcium carbonate shells. Although our monitoring of these chemical changes has improved, we have not developed effective tools to translate observations, which are typically of the pH and carbonate saturation state, into ecologically relevant predictions of biological risks. One potential solution is to develop bioindicators: biological variables with a clear relationship to environmental risk factors that can be used for assessment and management. Thecosomatous pteropods, a group of pelagic shelled marine gastropods, whose biological responses to CO_2_ have been suggested as potential bioindicators of OA owing to their sensitivity to acidification in both laboratory and the natural environment. Using five CO_2_ exposure experiments, occurring across 4 seasons and running for up to 15 days, we describe a consistent relationship between saturation state, shell transparency, and duration of exposure, as well as identify a suite of genes that could be used for biological monitoring. We clarify variations in thecosome responses due to seasonality, resolving prior uncertainties and demonstrating the range of their phenotypic plasticity. These biomarkers of acidification stress can be implemented into ecosystem models and monitoring programs in regions where pteropods are found, while the approach will serve as an example for other regions on how to bridge the gap between point-based chemical monitoring and biologically relevant assessments of ecosystem health.

**Summary Statement:** Despite seasonal variability, pteropods exposed to acidification over multiple seasons reveal consistent patterns in gene expression and shell condition that can be used as bioindicators of ocean acidification stress.

## Introduction

Anthropogenic activity since the industrial revolution has released a large amount of carbon dioxide (CO_2_) into the atmosphere (~2400 GtCO_2_)(Friedlingstein et al., 2022; IPCC, 2022), a substantial fraction of which (~30%) has been dissolved in ocean waters(Gruber et al., 2019). The addition of this excess CO_2_ into marine systems profoundly modifies the acid-base chemistry of seawater, shifting equilibrium towards a lower pH and a lower saturation state for calcium carbonate compounds (Doney et al., 2009). This process, called ocean acidification (OA), thus tends to dissolve the skeletal components of marine organisms made from calcium carbonate, and has been shown to impact rates of growth, calcification, gene expression, survival, and development for a range of species (Espinel-Velasco et al., 2018; Kroeker et al., 2010), although sensitivity is taxonomically heterogeneous, often relating to the presence of a shell, it is additionally moderated by ecological, physiological, and ontogenetic factors (Hancock et al., 2020; Leung et al., 2022).

As our awareness of the repercussions of acidification has risen, there have been concerted efforts to increase monitoring of the carbonate chemistry and acidification of both open-ocean and coastal systems (Bates et al., 2014; Tilbrook et al., 2019; Wang et al., 2013). The objective of monitoring programs is to synthesize observations of chemistry and biology into information relevant for policy development and resource management (Cross et al., 2019; Doney et al., 2020; McLaughlin et al., 2015). Although there has been progress in implementing observations of chemistry into regional predictive assessments of ecosystem risk (Cooley et al., 2015; Hare et al., 2016), translating laboratory observations of organismal OA sensitivity to in situ impacts has been difficult to accomplish (Doo et al., 2020; Weisberg et al., 2016). Plasticity in species responses, as well as seasonal or regional variability of exposure, parental provisioning, ontogeny, and other environmental and biological factors, play large roles in our current uncertainty of population level effects of OA.

Bioindicators are biological variables with a clear relationship to environmental risk factors that can be applied to implement ecologically relevant water quality criteria and model risk thresholds for assessment and management (Bednarsek et al., 2019; Weisberg et al., 2016). One common example is the abundance of coliform bacteria as a proxy for fecal contamination in water. In the case of OA, bioindicators are typically compared with chemical measures of acidification, such as the saturation state of calcium carbonate compounds used for calcification. These organismal indicators of ecological state hold promise for improved OA monitoring, as these phenotypic presentations of health are typically based on responses integrating over longer timescales and broader spatial scales than the point-based chemical measurements, while also being directly linked to the metrics stakeholders care about – the health of organisms and ecosystems.

One taxonomic group that has emerged as a potentially providing useful bioindicators of ecosystem acidification stress is the aragonite-shelled euthecosomatous pteropods (often referred to as thecosome pteropods, or just pteropods in the literature, and commonly referred to as sea butterflies). The respiration rate, gene expression, and shell condition of these organisms are influenced by OA in the laboratory (Bednarsek et al., 2019; Maas et al., 2018; Manno et al., 2017), and there is strong evidence that their shell condition can be an indicator of OA exposure in modern oceanic conditions (Bednaršek et al., 2022; Bednarsek and Ohman, 2015; Bednaršek et al., 2012; Mekkes et al., 2021; Niemi et al., 2021). Pteropod shells are made of a more soluble calcium carbonate compound (aragonite) than the form used by most planktonic species, and are impacted at a higher saturation state (ΩAr =1.5) than would be predicted from pure chemical equilibrium (ΩAr =1.0), the point at which aragonite is predicted to dissolve (Bednarsek et al., 2019), suggesting that their shell condition could serve as an “early warning” of OA impacts for other shelled species. Natural phenology and seasonal cycles have, however, been demonstrated to influence phenotypic responses of thecosomes to OA (León et al., 2020; Maas et al., 2020), confounding and potentially obscuring the correlations between saturation state and the gene expression, respiration, and shell condition of these potential bioindicators. A metanalysis of the response metrics was only able to find consensus on thresholds of response (Bednarsek et al., 2019), rather than develop consistent predictive relationships between saturation state and a response variable.

Previously proposed pteropod-based bioindicators that relate saturation state to biological indicator include shell dissolution (using SEM), shell calcification (using calcein staining), and survival (Bednaršek et al., 2017). Both calcification and survival are only valuable for the detection of tipping points, as they require the capture and maintenance of organisms for a period of time to assess the biological response and can’t be used with wild caught animals to assess their prior exposure. Shell dissolution using SEM evaluation of mild and/or severe dissolution, by contrast, can be applied to animals collected directly from the wild. Studies have shown that SEM evaluation, which is qualitative rather than quantitative, is not a sensitive metric across the full range of saturation states experienced by thecosomes, and results are not highly repeatable between users (Oakes et al., 2019). Furthermore, there is an extensive debate in the literature as to the effects of animal handling, associated with the lab studies that use the SEM method (Bednarsek et al., 2016; Miller et al., 2023; Peck et al., 2016a; Peck et al., 2016b), making it a less desirable approach to widely implement in monitoring programs. SEM is only one of the various shell quality metrics that have been implemented, however, with various other analyses, including transparency, opacity, and a Limacina Dissolution Index also used throughout the literature (reviewed in Oakes et al., 2019). Of these the quantitative microscopy techniques (dry shell opacity and transparency) have been revealed to have both high repeatability and sensitivity across a large range of saturation states (Bergan, 2017; Oakes et al., 2019). The work has not been done yet to demonstrate quantitative relationships between duration and severity of exposure, or to determine whether there are seasonal or ontogenetic differences in CO_2_ sensitivity.

Beyond shell condition, respiration rate and gene expression have been suggested as potential bioindicators for pteropod OA response. Respiration rate response to acidification has been shown to be variable and dependent on the presence of co-stressors (Comeau et al., 2010; Lischka and Riebesell, 2012; Seibel et al., 2012), although the cause of this variability is unclear. Gene expression patterns are emerging that suggest similar suites of upregulated and downregulated genes are present during periods of high CO_2_ exposure across species and studies (Johnson and Hofmann, 2017; Maas et al., 2018; Maas et al., 2020). They are overlaid by patterns of seasonal gene expression that could be either signals of environmental exposure or ontogenetic and developmental phenology that have, thus far, made it difficult to isolate distinct biomarkers that are associated with in situ exposure to CO_2_. To precisely identify biomarkers we thus require studies that disentangle seasonal responses from exposure level responses, isolating natural variability from CO_2_ specific responses. The best experimental design is thus a repeated controlled laboratory experimental design embedded within natural seasonal variability.

*Limacina retroversa* is a dominant euthecosome pteropod species in the temperate North Atlantic, broadly distributed in coastal and open-ocean waters (Bé and Gilmer, 1977). Although pteropods, with their delicate shells and mucous-web feeding, are notoriously difficult to culture (Howes et al., 2014), substantial progress has been made in rearing *L. retroversa* in aquaria, making it a strong candidate as a model pteropod species (Bergan et al., 2017; Maas et al., 2018; Thabet et al., 2015). This study is designed to assess seasonal patterns of physiological responses in the thecosome pteropod, *L. retroversa*, to laboratory exposure to CO_2_. These organisms are found throughout the year in the Gulf of Maine region, where they experience natural seasonal cycles in saturation state (Maas et al., 2020; Wang et al., 2017). Prior work in the area has documented pulses of reproduction in the spring and fall, reduced shell transparency in the winter when environmental CO_2_ is naturally elevated, increased respiration rate during the spring bloom, and seasonally distinct patterns of gene expression (Maas et al., 2020). Additionally, laboratory experiments conducted exclusively during the spring bloom have demonstrated that there are increases in respiration rate, reduced shell transparency, and increasing numbers of differentially expressed genes with increasing intensity and duration of CO_2_ exposure (Maas et al., 2018). Here we use four seasonally repeated laboratory exposures to CO_2_ during 2014 to determine whether there is plasticity in such responses in relation to ontogenetic and environmental variation, while identifying consistent biological markers of acidification stress that would be appropriate for implementation into biomonitoring projects (Fig. 1). A fifth experiment in spring of 2015 was additionally conducted to expand shell response data.

**Figure 1:**
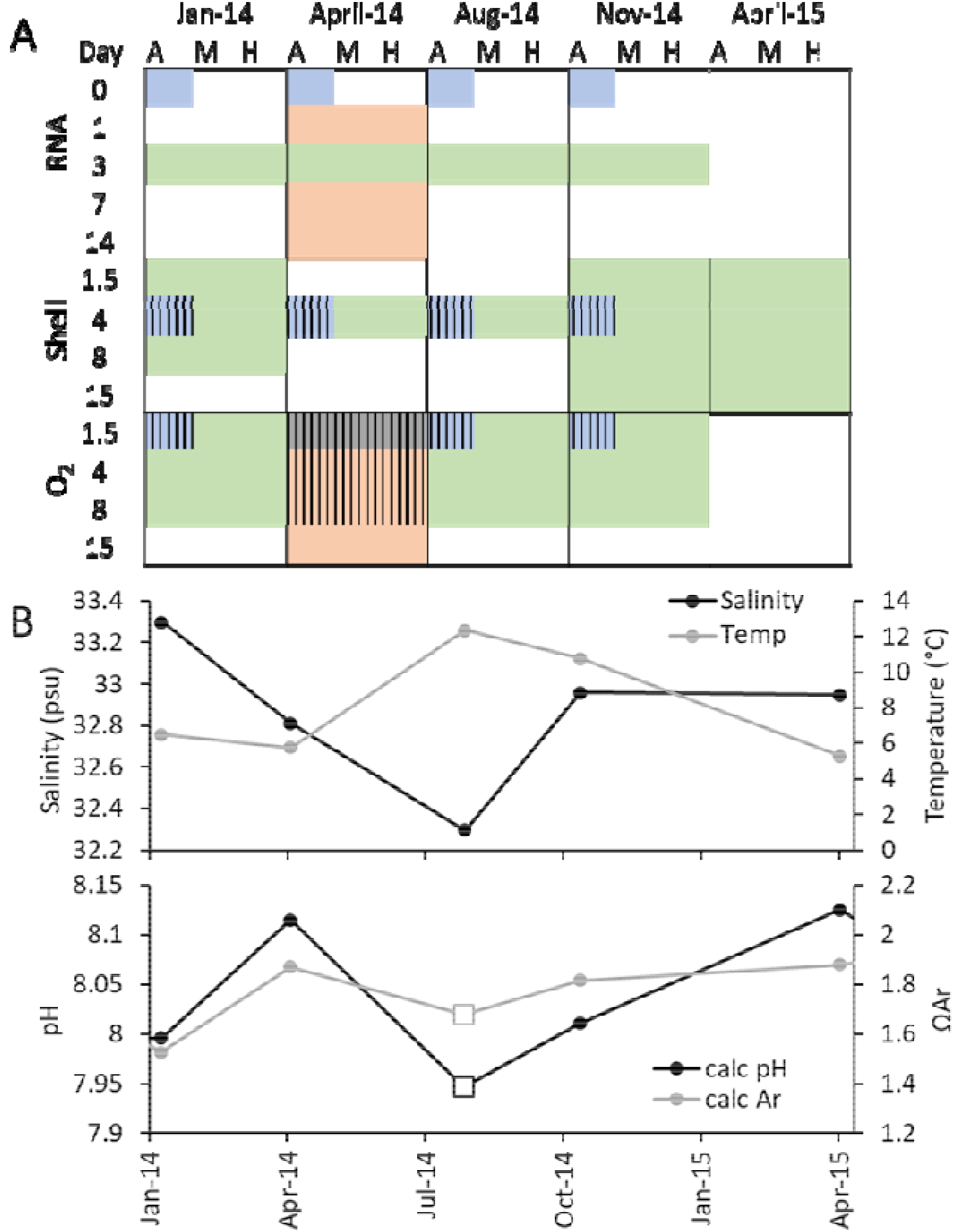
Sampling design and seasonal profiles. The sampling plan (A) consisted of taking animals from five different cruises and exposing them for up to 15 days to three CO_2_ treatment levels: ambient (A), medium (M), and high (H). Detailed carbonate chemistry from these exposures is available in Table 1. Some data from these experiments has previously been published in a durational study (Maas et al. 2018; peach), and an in situ seasonality study (Maas et al. 2020; blue), but the majority of the samples described herein were analyzed explicitly for this project (green). Samples from a prior analysis that were also used in this analysis are demarcated by stripes. (B) Upper water column hydrographic (CTD; average 0-60 m) and carbonate chemistry bottle sampling (bottle sampling; average 0-60 m, n=3) demonstrates the seasonal variability in environmental conditions which are associated with phenological patterns of growth and reproduction, with a pronounced spring spawning event peaking around May and a smaller late fall reproductive event as described in Maas et al. (2020).

**Table 1:** Average carbonate chemistry parameters from seasonal laboratory exposures. Dissolved Inorganic Carbon (DIC), Total Alkalinity (TA), and pH (total scale) were each measured in all three treatments: ambient, medium, and high (A, M, H, respectively; sampling frequency and full carbonate chemistry in Table S2). The DIC/TA pairs were used with measured salinity and temperature (Temp) values to calculate pCO_2_ (ppm ± standard error) and the aragonite saturation state (ΩAr ± standard error) using the program CO2SYS.

| Sampling period | Treat. | Salinity | Temp (°C) | DIC ($\mu\text{mol kg}^{-1}$) | TA ($\mu\text{mol kg}^{-1}$) | pH | $p\text{CO}_2$ | $\Omega\text{Ar}$ |
| --- | --- | --- | --- | --- | --- | --- | --- | --- |
| Jan 2014 | A | 32 | 8.02 | 2101.1 | 2248.2 | 7.95 | 421 ± 14 | 1.69 ± 0.05 |
| | M | 32 | 8.02 | 2177.8 | 2250.0 | 7.73 | 728 ± 13 | 1.08 ± 0.02 |
| | H | 32 | 8.02 | 2224.6 | 2257.2 | 7.59 | 1016 ± 32 | 0.81 ± 0.03 |
| April 2014 | A | 34 | 8.05 | 2084.7 | 2219.0 | 7.97 | 471 ± 11 | 1.54 ± 0.03 |
| | M | 34 | 8.05 | 2167.0 | 2223.4 | 7.74 | 852 ± 20 | 0.95 ± 0.02 |
| | H | 34 | 8.05 | 2202.8 | 2219.2 | 7.60 | 1189 ± 45 | 0.71 ± 0.02 |
| Aug 2014 | A | 34 | 7.77 | 2039.8 | 2182.5 | 8.02 | 430 ± 14 | 1.60 ± 0.04 |
| | M | 34 | 7.77 | 2113.4 | 2188.3 | 7.75 | 718 ± 40 | 1.07 ± 0.06 |
| | H | 34 | 7.77 | 2154.0 | 2189.7 | 7.64 | 985 ± 39 | 0.80 ± 0.03 |
| Nov 2014 | A | 33 | 7.57 | 2084.6 | 2210.7 | 8.01 | 476 ± 17 | 1.48 ± 0.03 |
| | M | 33 | 7.57 | 2179.2 | 2211.9 | 7.71 | 1019 ± 100 | 0.81 ± 0.06 |
| | H | 33 | 7.57 | 2209.4 | 2215.4 | 7.58 | 1258 ± 72 | 0.65 ± 0.04 |
| April 2015 | A | 33 | 8.04 | 2081.3 | 2218.5 | 7.99 | 449 ± 3 | 1.58 ± 0.01 |
| | M | 33 | 8.04 | 2152.3 | 2221.8 | 7.78 | 750 ± 35 | 1.04 ± 0.04 |
| | H | 33 | 8.04 | 2202.0 | 2216.8 | 7.59 | 1183 ± 61 | 0.70 ± 0.04 |

## Methods

*Limacina retroversa* were collected at five different times from the Gulf of Maine, and held in three CO_2_ treatments for up to 15 days of exposure. They were sampled for respiration rate, shell condition and gene expression at various periods throughout their exposure (Fig. 1). The hydrography of the seasonal sampling, as well as the general abundance and vertical distribution of the organisms during these seasonal cruises has been reported previously (Maas et al., 2020; Wang et al., 2017), providing the ecological context of the seasonality.

### Seawater and Animal Sampling

Adult *Limacina retroversa* were collected for physiology experiments during short (1-3 day duration) seasonal cruises conducted within the Gulf of Maine (42° 22.1’ - 42° 0.0’ N and 69° 42.6’ - 70° 15.4’ W) beginning January 29^th^, April 25^th^, August 19^th^ and November 4^th^ 2014 from aboard the *R/V Tioga*, as described in Maas et al (2018; 2020). There was an additional cruise in April 25^th^ of 2015 where animals were retained exclusively for studies of the shell as described in Bergan et al (2017). Prior to animal capture, water for animal maintenance was pumped from ~30 m depth using a submersible pump and filtered using a 63 µm mesh filter. Some of this water was transferred into 1 L jars that were placed in a refrigerator at 8 °C to keep it at temperature for animal transport. The rest of the water was stored for transport in large covered plastic bins. On the first day of each cruise this water was transported to an 8 °C walk-in environmental chamber at Woods Hole Oceanographic Institution where it was deposited into a holding tank (>400 l) and recirculated past a 1-µm filter throughout the duration of experiments (maximally 2 weeks).

*L. retroversa* were sampled using a Reeve net with 333-µm mesh and a large cod end. These were short duration (<1 h) oblique tows (vertical speed 5-10 m min^-1^, 1-2 knot vessel speed) through depth strata where densities had been revealed to be high via prior MOCNESS (Multiple Opening and Closing Net and Environmental Sampling System) and seasonal sampling (generally 80-50 m in the offshore stations and 50-25 m at nearshore sites; described in Maas et al. 2020). Once the net was onboard, the cod end was promptly divided among several buckets and diluted. *L. retroversa* were sorted from other taxa using a wide bore plastic pipette and placed at densities between 20-40 ind. L^-1^ in the 1 L jars with refrigerated pre-filtered *in situ* water. These jars were stored in coolers or the 8 °C refrigerator for transport back to the lab.

### Experimental Exposure to CO_2_

As detailed in Maas et al. (2018), in-situ collected pre-filtered water was transferred into three ~100 L pre-equilibration tanks and allowed to bubble for ~12 h prior to the introduction of animals. Gas concentrations were generated with mass flow controllers (Aalborg, Orangeburg, NY, USA) that combined local compressed ambient air (380 to 440 µatm) and CO_2_ to achieve a medium (~800 ppm) and high (~1200 ppm) treatment (Table 1) as described in White et al. (2013). These treatments were chosen to yield supersaturated, near-saturated, and undersaturated conditions in each season, with some variability owing to seasonal differences in ambient conditions; these three levels of aragonite saturation state are referred to as CO_2_ treatments, while the precise saturation state is referred to as the intensity of the exposure in subsequent analyses. When organisms were brought to the lab after the cruise, this pre-bubbled water was pumped into three 12-L glass experimental carboys per treatment (a total of 9 carboys placed in a semi-randomized pattern in the environmental chamber), where bubbling was continued using micro-bubbling stones.

Animals were individually distributed randomly into the pre-bubbled experimental carboys at densities of 20-25 individuals L^-1^. In all of the experiments except November 2014, only those individuals that had been collected on the last day of the cruise were used in the experiments. Due to low sampling density in November, individuals were used from the last two days of cruise sampling. After placement into the experimental containers, animals were fed a mixture of *Rhodomonas lens* and *Heterocapsa triquetra*, and this feeding regime was repeated once every 2 or 3 days as detailed in Thabet et al. (2015) and Maas et al. (2018). Water changes were conducted after one week of captivity using the remaining water in the holding tank following the protocol of pre-bubbling as mentioned above.

### Carbonate Chemistry Analyses

The temperature of the environmental chamber was measured continuously throughout the experiment using the temperature sensor associated with the FireSting fiber-optic oxygen meter (PyroScience, Aachen Germany). Salinity was measured from the experimental carboys using a seawater refractometer (Hanna Instruments, model 96822) every 2-3 days and during water changes. The pH of each carboy was determined using a USB 4000 spectrometer with an Ls-1 light source and a FIA-Z-SMA-PEEK 100 mm flow cell (Ocean Optics, Dunedin, FL, USA) every 2-3 days using a 2mM m-Cresol purple indicator dye and as described in Maas et al. (2018). Measurements of dissolved inorganic carbon (DIC) and total alkalinity (TA) were conducted on bottle samples collected from pre-bubbled water at the start of the experiment, before and after the water change, and the end of the experiment as detailed in Maas et al. (2018). Samples were collected in 250mL Pyrex borosilicate glass bottles, each of which was poisoned with saturated mercuric chloride, following published best practices for seawater CO_2_ measurements (Dickson et al., 2007; Riebesell et al., 2010). DIC was measured using an Apollo SciTech DIC auto-analyzer, while TA was measured using an Apollo SciTech alkalinity auto-titrator, a Ross combination pH electrode, and a pH meter (ORION 3 Star) based on a modified Gran titration method (Wang et al., 2013). The aragonite saturation state (ΩAr) and pCO_2_ during each experimental time point were calculated using concurrently collected DIC-TA pairs, or using the more frequent pH measurements with the TA pair from the closest water change. The seawater carbonate chemistry calculations were made with the CO2SYS software (Pierrot et al., 2006), using constants K1 and K2 by Lueker et al. (2000), the KHSO_4_ dissociation constant from Dickson (1990), and the borate relationship from Lee et al. (2010).

### Respiration Experiments

Respiration measurements were started on days 1, 3, and 7 of the seasonal experiments, following the protocol detailed in Maas et al. (2018). Prior to being placed in respiration chambers, individuals were removed from the experimental carboys and placed in a 1-L beaker at densities of 15 ind L^-1^ with 0.2 µm filtered pre-bubbled in situ treatment-specific water for 8-12 h to allow for gut clearance. Then they were transferred to custom small volume glass respiration vials containing fresh 0.2 µm filtered pre-bubbled in situ treatment specific water. Each vial contained an optically sensitive “spot” (OXFOIL: PyroScience, Aachen Germany) for oxygen sensing. The volume of the chamber was then adjusted to between 2-3 mL and closed. A control, filled with water but left without an animal, was set up every fourth chamber for bacterial respiration measurements. The oxygen concentration in each chamber was then measured using a FireSting fiber-optic oxygen meter (PyroScience, Aachen Germany). The optode was calibrated using air saturated seawater and zeroed using a 2% sodium sulfite solution at the start of each seasonal experiment. At the conclusion of the respiration experiment (~24 h) the O_2_ concentration was again measured for each chamber. Consequently, animals were sampled after a total exposure duration of 36 h, 4 d, and 8 d. The chambers were then weighed wet and emptied and weighed dry to get an accurate estimate of the water volume. Each organism was visually inspected to verify if it was still alive and then was briefly rinsed with DI water, placed in pre-weighed aluminum dish, and weighed on a Cahn microbalance (C-33). After weighing, individuals were put in a drying oven at 70 °C for > 3 days and weighed again. Final oxygen consumption rates were calculated based on the wet weight and the change in oxygen consumption between the final and initial oxygen measurements (µmol O_2_ g^-1^ h^-1^). The results were not corrected for the low bacterial respiration from the control chambers which averaged to 0.0002 µmol O_2_ h^-1^, which accounts for less than 5% of the oxygen consumption in an experiment.

There was some variability in the temperature of the environmental rooms across the experiments. Although the rooms achieved 8.1 ± 0.5°C for most experiments, an equipment failure for the chiller unit during the first day of one cruise (August 2014; 5.6°C) resulted in unexpectedly cold temperatures and the need for a temperature correction across the dataset. The average temperature of the 24 h respiration experiment was used for the original temperature (T_i_) and the adjusted rates (R_f_) were calculated at 8.0°C using a temperature coefficient (Q_10_) of 2 according to the equation:

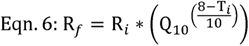

where R_i_ is the final oxygen consumption rate measured for each individual. Although Q_10_ is known to be species-specific in pteropods (Seibel et al., 2007), there are no published studies of Q_10_ for *L. retroversa*. This chosen coefficient is mid-range for the published Q_10_ of congeners (1.6-2.3; Ikeda, 2014; Maas et al., 2011). All temperatures experienced by the pteropods were within the normal annual range of their natural environment (Maas et al., 2020), and the total variation in the average temperature among the experiments was small (−2.4 to + 0.6 °C), so slight variations in actual Q_10_ would not substantially influence the calculated temperature-corrected respiration rate.

Statistical analysis involved first testing for an effect of season, CO_2_ treatment, and duration of exposure on the log of mass-specific respiration across the full dataset using a GLM, with log of wet mass as a covariate and using the statistical program SPSS. Each season was then explored separately to determine the effect of CO_2_ treatment severity and duration of exposure and differences explored using bonferonii post hoc tests. Metabolic rate was additionally analyzed based on the intensity of the CO_2_ exposure level.

### Shell Transparency Analyses

Shell transparency analysis was conducted on individuals from all five of the seasonal cruises after 4 days of exposure. For all of the 2014 cruises, individuals that had been used in the respiration experiments were dried as described above, and then analyzed for shell transparency following the methods of Maas *et al*. and Bergan *et al*. (2017; Maas et al., 2020). For the 2015 cruise, animals were removed directly from the experimental treatments for transparency analysis. Additional samples from January 2014, November 2014 and April 2015 were taken from individuals exposed for a duration of 36 h, 4 d, 8 d, and 15 d and were used to explore general patterns of response to intensity and duration of CO_2_ exposure.

To remove body tissue, individuals were placed in 8.25% hypochlorite bleach for 24-48 h, rinsed in DI water, and dried. These individuals were then photographed under a stereoscopic light microscope at 25x magnification with a background greyscale value of 255. Images were analyzed by thresholding the image to black and white and the aperture, as well as any holes, were manually cropped. The degree of light transmittance was then calculated using a custom MATLAB code with the mean greyscale value (range: 0–255) of the pixels of the shell divided by 255 to get a scale of 0 (black) to 1 (white). Using a GLM in SPSS we assessed whether there was an effect of season, treatment, and the interaction term on the transparency of the pteropod shells with differences explored using Bonferonii post hoc tests. We then used a GLM to determine whether there was a significant effect of season on the slope of the relationship between the log saturation state and log transparency for each duration of exposure.

A power equation relating saturation state and shell transparency was generated based on the average of the regressions from the 4 d experiments. To estimate the effect of duration, we applied this power equation to the 15-day shell transparency experiments. A theoretically “predicted” shell transparency was calculated for all shells from the medium and high treatments in these experiments, using equation 1 and the alpha value associated with the sampling period. The difference between the observed transparency and this “predicted” transparency was then plotted versus the duration of exposure to provide a regression that estimates the effect of duration on shell transparency after accounting for the effect of saturation state.

### Gene Expression

After 3 days of exposure, freely swimming pteropods were removed directly from the large carboys and immediately preserved in RNAlater for analysis of gene expression. Within each treatment, total RNA was extracted using the E.Z.N.A. Mollusc RNA Kit (Omega Biotek) from 5-6 biological replicates, each containing a pool of 5-9 pteropods. Three RNA samples were selected from each season and treatment combination (36 samples total) based on spectral profile and RNA yield, and these were submitted to the University of Rochester Genomics Research Center for sequencing. Libraries were constructed using TruSeq Reagents and then sequenced on 4 lanes of an Illumina HiSeq 2500 (9 samples per lane) as a High Output v4 125 bp PE project. The sequencing facility used Trimmomatic (v.0.32; Bolger et al., 2014) to eliminate adapter sequences (2:30:10) and removed low quality scores using a sliding window (4:20), trimming both trailing and leading sequences (13) and leaving only sequences with a minimum length of 25 for downstream use.

Reads from individual RNA samples were then aligned to the transcriptome that was previously generated and annotated in association with this project (Maas et al., 2018), allowing direct comparisons between studies. This assembly has been demonstrated to be sufficiently complete (BUSCO score C: 90.6% [S:65.1%, D:25.5%], F:8.0%, M:1.4%, n:978) to support the analysis (Simão et al., 2015). Alignment was completed using the pipeline packaged with Trinity (Haas et al., 2013), using Bowtie2 (v.2.2.3; Langmead and Salzberg, 2012), and estimates of abundances were made with RSEM (Li and Dewey, 2011). Read mapping statistics indicate a reasonable alignment rate of 73.8% ± 1.3% (Table S1). edgeR analysis of differential expression (DE) was performed with R v.3.0.1 (Robinson et al., 2010). Pairwise comparisons were made within each season between ambient samples and either medium or high samples to explicitly test for the effect of CO_2_ during each season. Genes were defined as DE if the false discovery rate and the p value were both < 0.05, and the log_2_-fold change was > 2 (corresponding to a four-fold change in expression).

To explore gross patterns of gene expression among days and treatments, TMM gene expression of all samples and treatments were log-transformed and then a Bray Curtis Similarity Matrix was generated for the data using PRIMER. Samples were plotted as an nMDS with the factors month and treatment to determine significant clustering. To explore the effect of CO_2_, a second nMDS was plotted using only those genes that were differentially expressed in a pairwise comparison. The environmental conditions from each experiment (Table 1) were then correlated with these patterns of gene expression to determine which factors were most predictive of transcriptomic responses. The DE genes were compared with prior analyses of the effect of the intensity and duration of CO_2_ exposure in the lab (Maas et al., 2018) as well as the prior *in situ* seasonal expression patterns (Maas et al., 2020). Finally, the log (x+1) TMM expression level of each gene was correlated with the saturation state of the sample to explore genes that may be valuable quantitative and consistent biomarkers of environmental exposure level.

## Results

### Carbonate Chemistry

During the five experiments, which ran for 15 days each, natural seasonal variability in seawater alkalinity and local CO_2_ levels translated into a range of different pH values and saturation states based on the bubbling of in-situ field collected seawater with ambient air (380 to 440 µatm) and CO_2_ to achieve medium (~800 ppm) and high (~1200 ppm) treatments (Table 1; Table S2). Although ambient treatment was always supersaturated (ΩAr =1.69-1.48), sometimes the medium and high treatments overlapped across seasons. Physiological response variables were thus analyzed using both the treatment level (as a nominal factor) and the saturation state (continuous variable).

### Respiration Experiments

Respiration rates, as measured after 36 h, 4 days, or 8 days of laboratory CO_2_ exposure during the four 2014 experiments, were significantly influenced by sampling period (F_3, 265_ = 32.615, p < 0.001) and treatment (F_2, 265_ = 3.481, p = 0.032), while duration of exposure was not significant (F_2, 265_ = 2.178, p = 0.115; Fig. 2A) in a full factorial analysis. Each sampling period was also independently analyzed for the influence of treatment level and duration. The only period where CO_2_ had an influence was April 2014, when level of CO_2_ treatment had a statistically significant effect (F_2, 68_ = 4.361, p = 0.017; Fig. 2B), and there was an interactive effect between CO_2_ and duration (F_2, 68_ = 2.655, p = 0.040). This was observed as an increasing difference between ambient and the other treatments as duration increased, and was supported by Bonferroni post hoc tests, which indicated that the medium and high treatments had a higher respiration rate than ambient after 8 days of exposure. The pattern is suggestive of a transition to an increased metabolic rate when exposed to CO_2_, reached earlier in the high treatment than the medium treatment in April.

**Figure 2:**
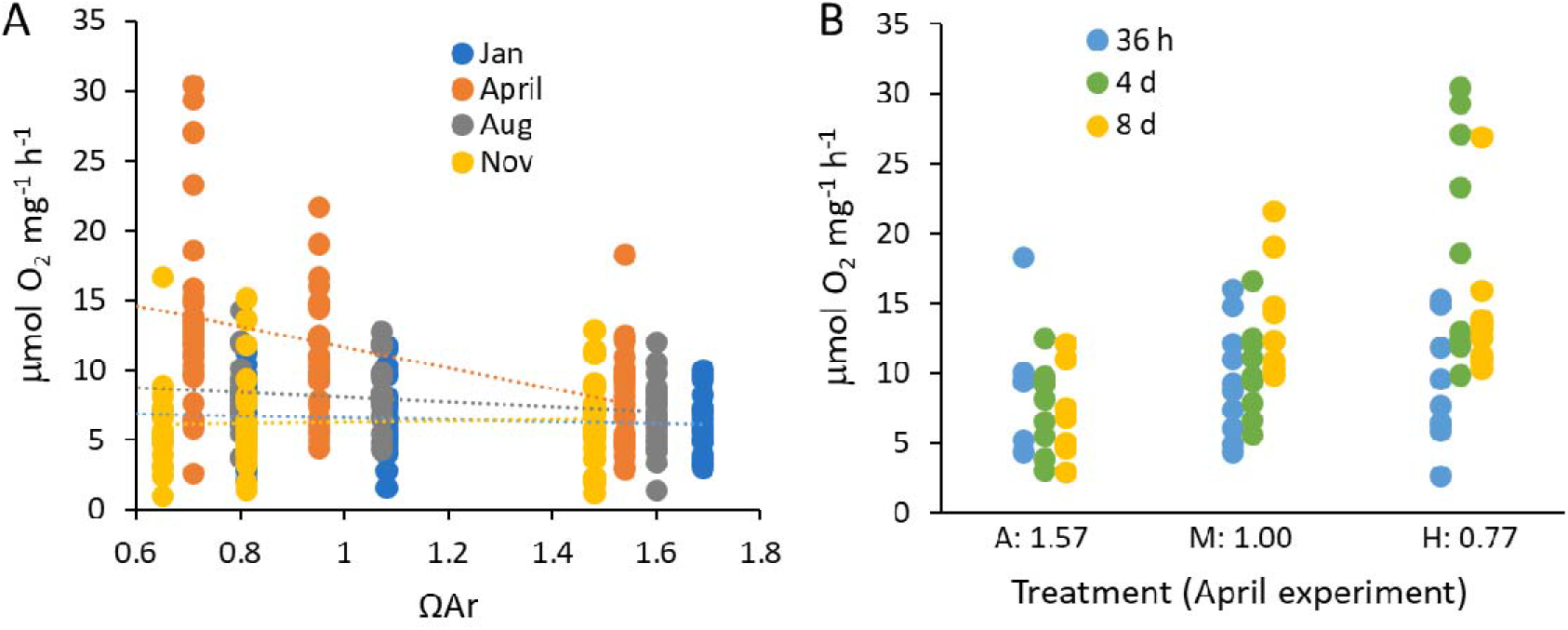
Measured mass-specific oxygen consumption compared with CO_2_ treatment levels. **A)** Metabolic rate for each sampling period (colors) is reported in comparison with the intensity of the CO_2_ exposure as indicated by measured aragonite saturation state. All exposure durations (36h, 4d, and 8d) are shown in aggregate as there was no overall effect of duration of treatment on metabolic rate for the full dataset. **B)** When seasons were analyzed independently, April was the only period where respiration rate was influenced by CO_2_, with a significant and interactive effect of both the duration (colors) and CO_2_ treatment level (x-axis).

### Shell Transparency

The shell transparency of organisms was measured for all 3 treatment levels after 4 days of laboratory exposure in each of the five sampling periods. Shells unexposed to elevated CO_2_ had higher transparency (max=0.94), while those exposed to higher intensity or longer duration CO_2_ exposure had a lower transparency (min=0.44). Although there was a significant effect of sampling period (GLM; F_4, 97_ = 63.451, p < 0.001) and a clear effect of treatment (F_2, 97_ = 44.441, p < 0.001), there was no interactive effect between sampling period and treatment (F_8, 97_ = 1.684, p = 0.112), meaning that there was no significant difference in the relationship between shell transparency and treatment among sampling periods (Fig. 3A). Our earlier work in the Gulf of Maine region has demonstrated lower transparency in shells from field caught pteropods during the winter, when saturation state is naturally at its lowest point (Maas et al., 2020). Shells from January retained this significantly lower transparency compared to all of the other seasons, while those from April 2014 and November had intermediate transparency, and those from August and April 2015 had the highest transparency (Bonferonii post hoc analysis; Fig. 3A). Synthesizing this data provides a quantitative relationship between the saturation state and shell transparency that is best expressed as a power function:

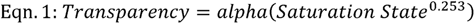

where the exponent is calculated as the average exponent from each independent season (n=5; standard error = 0.032). The constant (*alpha*) is related to the prior exposure to CO_2_ in the environment, and was strongly correlated (R² = 0.9847) with the previously measured shell transparency of field caught organisms (Day 0) from the same season (Maas et al., 2020):

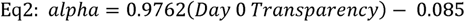

**Figure 3:**
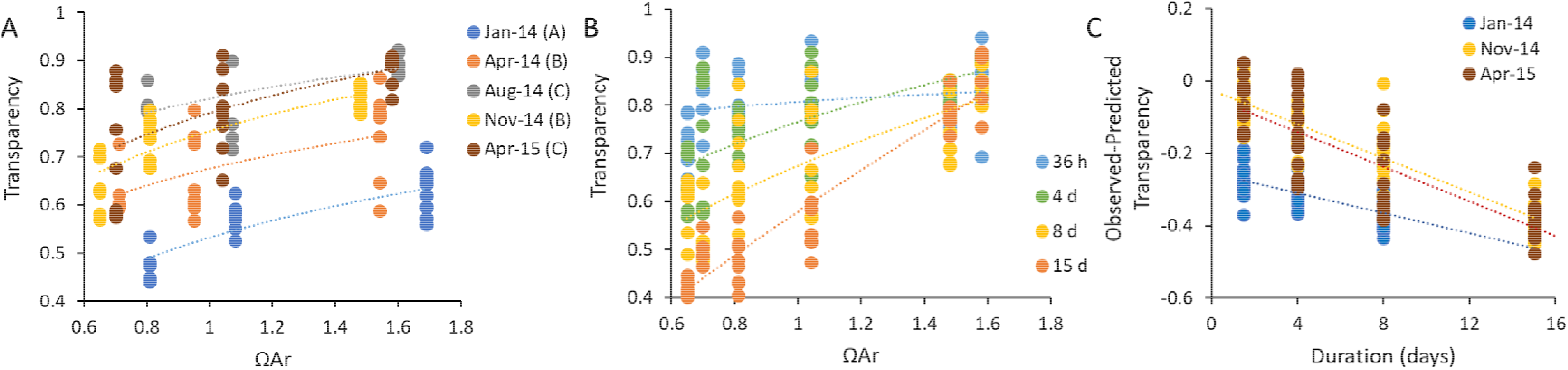
Effect of season, duration, and intensity on shell transparency. **A)**There was a consistent effect of the intensity of CO_2_ exposure (as indicated by aragonite saturation state) on shell transparency, where the month of capture (color; statistical grouping in parentheses) had a significant effect on the intercept (presumably ambient starting transparency; data shown after 4 days exposure, power regression), but no significant effect on the slope of the relationship between transparency and saturation state. **B)** The duration of exposure (color; time) had an interactive effect with the intensity of CO_2_ exposure, where increasing duration reduced the transparency at increasing intensity (shown excluding the January 2014 data points to provide a similar starting shell condition; power regression). **C)** Effect of the duration of exposure on shell transparency after observations were corrected for the intensity of CO_2_ exposure.

Using this equation with the highest observed shell transparence at Day 0, we estimate that alpha = 0.81 for pristine field caught shells uninfluenced by reduced environmental saturation states (~0.9).

To assess the combined effects of the intensity and duration of CO_2_ exposure on shell transparency, we conducted additional experiments during January 2014, November 2014, and April 2015 in which we used all three CO_2_ treatment levels, and measured transparency after 36 h, 4 d, 8 d and 15 d. Ambient treatments (ΩAr =1.69-1.48), retained a similar shell transparency for the duration of the experiment, while there was a statistically significant reduction in transparency over time for medium and high treatments, with an increase in the exponent of the relationship between saturation state and shell transparency over the duration of the experiments (Fig. 3B). Although there was a seasonal specific starting transparency that influenced the function, there typically was a statistically similar exponential relationship among seasons at each duration (SF1).

Since the effect of treatment did not statistically emerge until after the 36 h treatment (SF1), the effect of duration was modeled by first calculating a predicted day 4 transparency (D4T) for each observed saturation state in the duration experiments (using Eqn.1 and the observed alpha for each sampling period). The difference between the observed transparency and the D4T value can then be attributed to a change in shell condition caused by the duration of exposure (Fig 3C). There was no interactive effect between duration and season (F_5, 248_ = 1.885, p = 0.097). Knowing that the CO_2_ response is thought to have a threshold at ΩAr ~ 1.5 (Bednarsek et al., 2019), we tested the treatment levels separately. There was no effect of duration on shells from the ambient treatments (SF1; F_5, 76_=1.602, p=0.170), while there was a similar and significant effect of duration on both the medium (F_5, 75_=58.584, p<0.001) and high (F_5, 75_=52.090, p<0.001) treatments. Based on our November 2014 and April 2015 datasets we calculated this average linear relationship to be:

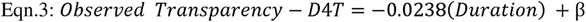

where beta is associated with the starting difference in shell transparency owing to seasonal variation, with an estimated beta of −0.0344 for a shell from a season with no prior exposure to OA. By setting the change in transparency to zero, we can calculate that the observed changes in shell transparency emerge after ~2.5 days of exposure (1.4 days prior to the day 4 measurements).

### Gene Expression

Our prior work on field caught *L. retroversa* demonstrated seasonal patterns of gene expression indicative of responses to *in situ* variation in saturation state (Maas et al., 2020). Additionally, a 14-day exposure study from April 2014 indicated an effect of both increasing CO_2_ intensity and duration on gene expression, but revealed an interactive effect between CO_2_ treatment and captivity duration (Maas et al., 2018). To minimize captivity effects and allow for a period of physiological response, samples here were analyzed in all four experiments from 2014 after 3 days of laboratory exposure to each treatment level. Patterns of total gene expression demonstrate a strong seasonality across the four sampling points studied, with January differing the most from the other three periods (Fig. 4A). When considering only the patterns of expression in transcripts that were statistically differentially expressed (DE) in at least one pairwise comparison between seasons, clustering was more strongly based on treatment (Fig. 4B). In this reduced dataset seasonal clustering was still present (particularly obvious in MD3; Fig. S2), emphasizing the strong seasonal influence on CO_2_ responsiveness.

**Figure 4:**
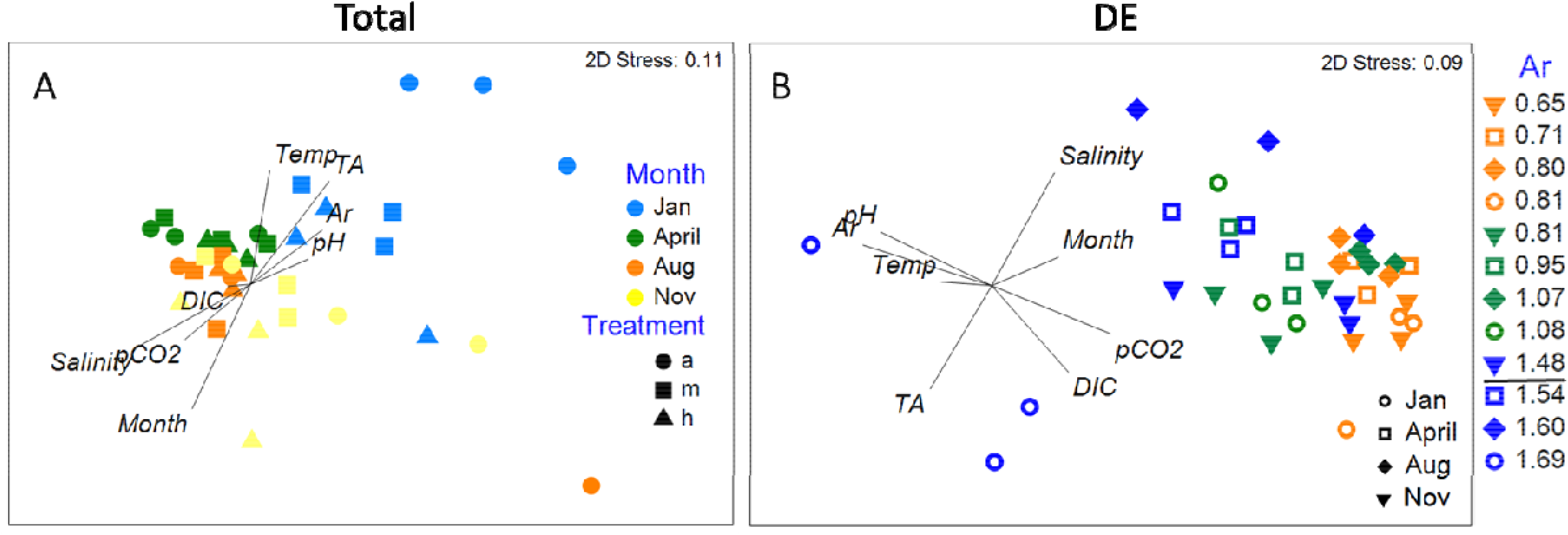
nMDS of total gene expression (A) and the expression of only those genes that were differentially expressed in one pairwise experimental comparison (B). Treatments are demarcated as ambient (a), medium (m), or high (h) in the left hand plot and by saturation state in the right hand plot with ambient (blue), medium (green), and high (orange) treatments falling at different levels owing to underlying differences in the carbonate chemistry. The theorized tipping point for saturation state (1.5) is noted in the legend. In the full comparison the experimental condition (Table 1) that was the best predictor of MDS1 (x-axis) was salinity, while the best predictor of MDS2 (y-axis) was month. In the analysis using only genes that were DE in one pairwise comparison, the experimental condition that was the best predictor of MDS1 was aragonite saturation state, while the best predictor of MDS2 was salinity.

Individuals in the January experiment, which experienced the lowest saturation states *in situ* prior to their capture, had the most pronounced response to elevated CO_2_ exposure (Table 2). The genes identified as DE in January shared the greatest number of DE genes with the previously published analysis of DE genes across seasons in freshly caught individuals in the field (Maas et al., 2020), emphasizing the similarity in response between field and laboratory exposures to CO_2_ (Table S3). April organisms were the next most responsive to laboratory exposure, while August, was the least (Table 2). The largest numbers of shared DE genes were in ambient versus high comparisons in January and April, followed by the medium versus high comparison from November (Table S3). Directionality of gene DE was generally conserved among studies and seasons (Table S3), emphasizing the reliability of these markers.

**Table 2:** Number of differentially expressed genes. Ambient (A), medium (M), and high (H), expression profiles were analyzed through pairwise comparisons within each month. DE genes from each month were then compared with those that were identified during a companion study of the effect of CO_2_ exposure duration (“duration” comparison), conducted during the April cruise and lasting for 14 days (Maas et al., 2018), and the number of shared genes is denoted. Finally, the DE genes from this study were compared with those that were differentially expressed during a seasonal study of in situ Limacina retroversa transcriptome expression (Maas et al., 2020) (“seasonality” comparison), and the number of shared genes are also indicated. Annotation of each DE gene, as well as the details of the expression patterns are noted in Table S3.

| comparison | Jan | Apr | Aug | Nov |
| --- | --- | --- | --- | --- |
| A vs H | 601 | 328 | 21 | 48 |
| A vs M | 300 | 11 | 44 | 7 |
| M vs H | 91 | 89 | 12 | 102 |
| Duration | 108 | 176 | 30 | 54 |
| Seasonality | 361 | 66 | 40 | 49 |

To search for potentially robust biomarkers associated with field and laboratory exposure to CO_2_, the list of genes identified in this study and our companion studies (Maas et al., 2018; Maas et al., 2020) were analyzed for shared DE genes (Table S3). Subunits of cytochrome c oxidase and NADH dehydrogenase were the most prominent components of the suite of genes typically expressed at higher levels (upregulated) in lower CO_2_ treatments or seasons with higher natural saturation state. Similarly, plasminogen, angiopoietin, fibrinogen, hemicentin, a zinc finger protein, and a number of unidentified sequences were generally expressed at higher levels (upregulated) in higher CO_2_ treatments or seasons with lower natural saturation state. These sequences were frequently annotated by Gene Ontology (GO) terms associated with the extracellular region and calcium ion binding. There were a number of genes (210) whose log (x+1) transformed TMM gene expression level had a high correlation (abs R>0.5) with saturation state (Fig. 5; Table S3). Generally, expression levels were, however, often variable and the coefficient of determination, which describes the proportion of variance explained by the correlation, was only high (R^2^ > 0.4) for a small subset of genes (30). The statistical correlation did not track with the frequency of the number of pairwise comparisons that were DE.

**Figure 5:**
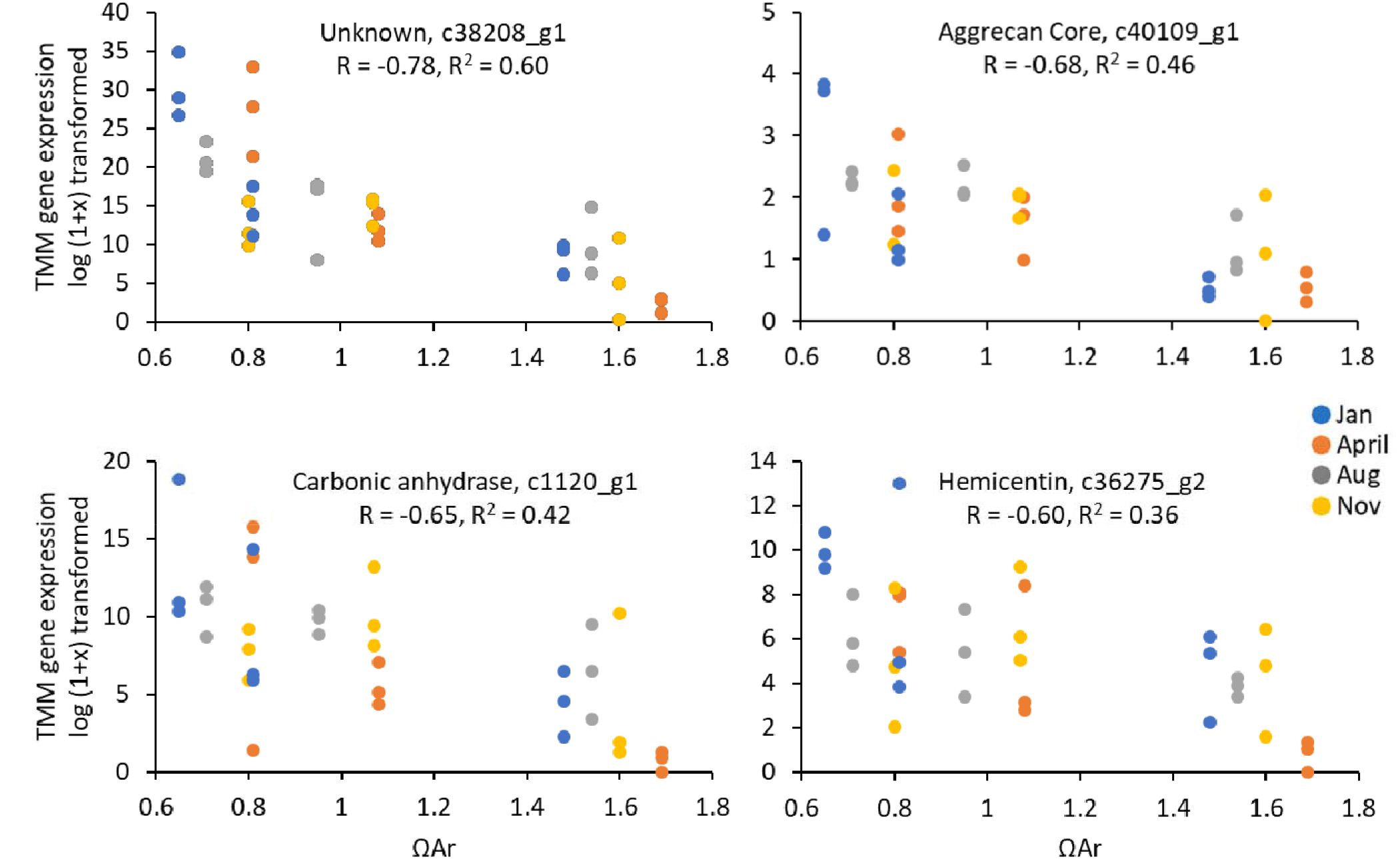
Effect of season and saturation state on a subset of genes. There was a relatively consistent effect of the intensity of CO_2_ exposure (as indicated by aragonite saturation state) despite the month of capture (color) on the gene expression (quantified as the TMM expression after log (1+x) transformation) for a number of genes after 3 days of exposure. Genes chosen for visual representation were not always the most highly correlated, but represent genes of interest from prior studies and DE analysis. A full list of the R and R^2^ for all DE genes are listed in TABLE S3.

## Discussion

Pteropod have emerged as a species whose responses to OA stress may serve as potentially valuable bioindicators, with studies reporting changes in pteropod shell condition, metabolic rate, and gene expression in response to CO_2_ exposure (Bednarsek et al., 2019; Bednarsek and Ohman, 2015; Bednaršek et al., 2012; Maas et al., 2018; Manno et al., 2017; Mekkes et al., 2021). To explore the scope of their seasonal phenotypic plasticity, and to identify consistent biomarkers of CO_2_ stress strongly correlated to saturation state, this study exposed *L. retroversa* to three treatment levels of CO_2_ for a duration of up to 15 days across five study periods spanning all four seasons. Owing to appreciable effects of natural seasonality, respiration was shown to be an unreliable marker for CO_2_ exposure as it was only associated with treatment during periods of high food availability. We identified, however, a suite of genes that may serve as appropriate biomarkers of OA stress after further studies that resolve the effect of duration of exposure. Finally, we demonstrated that shell transparency is consistently influenced by saturation state irrespective of season and, using these findings, we generated regressions of shell condition versus duration and intensity metrics. Together these results substantially advance our ability to quantitatively implement pteropod physiological responses as bioindicators of ecosystem OA exposure.

The metabolic rate of *L. retroversa* was previously demonstrated to vary seasonally in the wild, presumably in relation to food availability (Maas et al., 2020). This significant difference in metabolism, with higher rates of mass-specific oxygen consumption during the April spring bloom, had an interactive effect with CO_2_ exposure. In the present study, animals were fed identical rations during the experiments, but they would have experienced different food levels prior to collection that presumably affected their energy stores. We observed that metabolic rates were elevated in the more severe treatments (both by intensity and duration) in April and were not affected during other seasons. Our results indicate that higher food availability in conjunction with CO_2_ exposure is associated with an increased energetic expenditure in *L. retroversa* as measured by oxygen consumption. During the rest of the year, the animals appear to be using a basal metabolism, and may have no scope for increased energetic expenditures to apply to CO_2_ response. Similar seasonal variation in sensitivity to CO_2_ has been previously observed in silverside fish larvae (Baumann et al., 2018), where it was potentially attributed to maternal provisioning (Snyder et al., 2018). Food availability, which is related to body stores of energy, has consistently been demonstrated to influence CO_2_ sensitivity (Brown et al., 2018; Cominassi et al., 2020; Pansch et al., 2014; Seibel et al., 2012; Thomsen et al., 2013). Our dataset extends these findings by suggesting that responsiveness to CO_2_ must be inexorably linked with seasonal patterns of productivity and reproduction.

Shell condition, using a suite of different methodologies, has previously been shown to be impacted by CO_2_ gradients in natural populations in a variety of species of the pteropod family Limacinidae (Bednarsek and Ohman, 2015; Bednaršek et al., 2012; Maas et al., 2020; Mekkes et al., 2021; Niemi et al., 2021). Based on these previous *in situ* observations and a number of laboratory experiments, thresholds for changes in shell condition have been identified by meta-analysis near a saturation state of 1.5 (Bednarsek et al., 2019). Variations in species, ontogeny, experimental methodology, and treatment levels have, however, made it difficult to define a precise quantitative relationship between saturation state and shell response below that level. We have shown that despite the seasonally different starting levels (Maas et al., 2020), transparency was consistently influenced by the intensity of CO_2_ exposure in the laboratory in all 4-day seasonal experiments, and the threshold of an aragonite saturations state of ~1.5 was statistically upheld. The CO_2_ induced changes to transparency increased over time resulting in an interaction between intensity and duration of exposure, which can arguably be considered to be the total severity of exposure (Bednarsek et al., 2019). Our data demonstrates that changes in shell condition are negligible after 36 h of exposure, but emerge after ~2.3 days. There were very few individuals with transparency observed below the threshold of ~0.4, and these were only present in January in the 8 day high CO_2_ treatment, suggesting a lower biological limit. By measuring the transparency of a shell directly collected from the environment (T; which is the observed transparency) we can combine the equations 1 and 3 that relate transparency to saturation state, as well as the calculated alpha of 0.81 and beta of −0.0344, to model the range of durations (D), and intensities (ΩAr) experienced by the individual using the following equations:

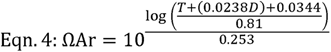

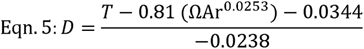

Based on these equations the transparency of shells gathered from the natural environment could provide a sense of the severity of ecological exposure and a range of intensities and durations of CO_2_ exposure organisms experienced in the period prior to capture (Fig. 6). These equations are easy to implement into ecosystem risk models, and the low cost and simplicity of the measure makes it attractive for implementation into general ecological monitoring projects.

**Figure 6:**
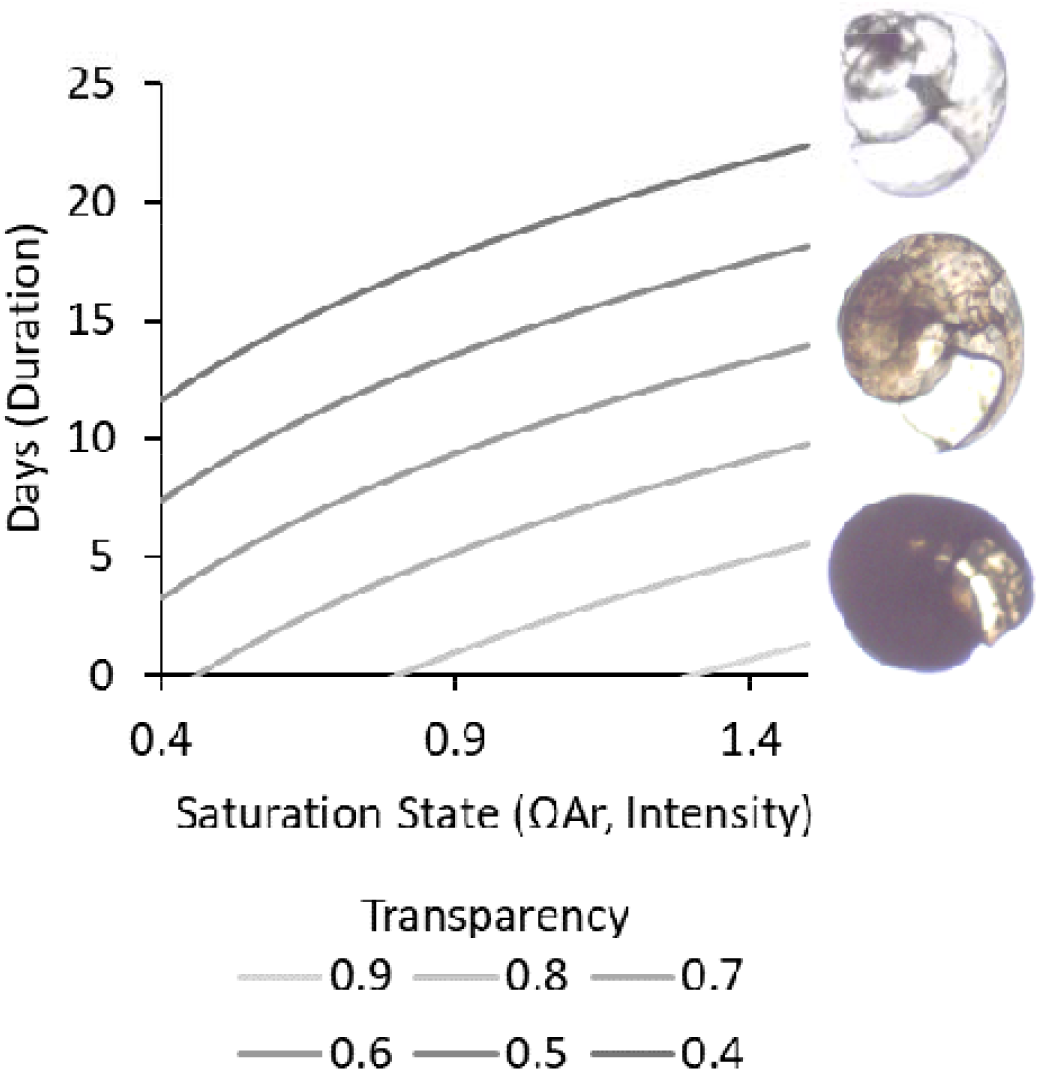
Potential environmental exposure based on observed shell transparency. Given a specified shell transparency measured from wild-caught organisms (lines), the ranges of intensity (x-axis) and duration (y-axis) required to produce the observed severity of effect can be calculated between the observed transparency range of 0.9-0.4 and the hypothesized tipping point of ΩAr =1.5 after more than two days of CO_2_ exposure.

Prior analyses of gene expression from freshly caught organisms emphasized a strong seasonal pattern in physiology reflecting annual responses to carbonate chemistry, food availability, and ontogenetic/developmental cycles (Maas et al., 2020; Wang et al., 2017). In this study, despite animals being held in similar temperature, carbonate chemistry, and food conditions, seasonality was still apparent after three days of captivity. This seasonal pattern of gene expression was apparent even when only considering the genes that were differentially expressed in response to CO_2_. Specifically, organisms from January, the period when the natural environment was measured to have the lowest chlorophyll *a* and lowest saturation state of all sampling points (Maas et al., 2020), had the highest transcriptomic responsiveness to CO_2_. April, the period of highest reproduction and the spring bloom, had the next highest number of DE genes in response to laboratory CO_2_ exposure, although the response was mainly measured in the high CO_2_ treatment. These results verify that these animals are actively responding to shifts in CO_2_, and suggest that this responsiveness has either an ontogenetic, nutritional, or prior-exposure dependence. Interpreting this responsiveness in the context of “sensitivity” or fitness is not possible from our data.

Despite the seasonal variability in gene expression response to laboratory CO_2_ exposure we identified a suite of genes that were frequently at a lower expression in the higher CO_2_ treatments. Those that were identified were associated with energetic metabolism – specifically cytochrome oxidase subunits and NADH dehydrogenase subunits. This is consistent with prior documented transcriptomic responses to high CO_2_ in other thecosome pteropods (Maas et al., 2018; Maas et al., 2015; Moya et al., 2016), urchins (O’Donnell et al., 2010; Todgham and Hofmann, 2009), and coral (Moya et al., 2012). These transcripts may, however, be poor biomarkers of OA exposure owing to the lack of specificity of their response (they have been identified as biomarkers of anoxia and heat stress in other invertebrates (Ma and Haddad, 1997; Mahadav et al., 2009; Woo et al., 2013)). The correlation between the log (x+1) TMM gene expression levels and the saturation state of the sample were low and the expression levels highly variable for these genes (R = 0.01-0.44; R^2^ < 0.19), which would prevent them from serving as reliable indicators of exposure to acidified water.

In the elevated CO_2_ treatments there were many transcripts with Gene Ontology (GO) terms associated with the extracellular region and calcium ion binding that were upregulated in medium and high treatments. Additionally, specific genes were consistently DE in response to higher CO_2_ including plasminogen, angiopoietin, fibrinogen, hemicentin, a zinc finger protein, and a number of unannotated sequences (Table S3). Most of these were also flagged as being DE in the durational study (Maas et al., 2018; Fig. 1) and were up-regulated in January during the *in situ* seasonal analysis of gene expression (Maas et al., 2020; Fig. 1). Some have similarly been identified as being DE in studies of other species of thecosome pteropods (Koh et al., 2015; Moya et al., 2016) and the much more distantly related shelled heteropods (Ramos-Silva et al., 2022). Many of these genes appear, however, to be influenced by seasonality; although the overall pattern of expression was consistent, the precise gene expression level in relation to saturation state was variable, leading to lower correlations and coefficients of determination. This indicates that although these DE genes are consistently effected by OA, there is background variability in their expression levels that are indicative of either pre-exposure in the environment, or seasonal and ontogenetic differences in expression level. Although understanding the processes controlled by these transcripts are important for our understanding of OA, these genes are thus not be ideal biomarkers of pteropod exposure to changing saturation state.

There were, however, 210 genes where the correlation between the log (x+1) TMM gene expression levels and the saturation state of the sample was relative high (abs R> 0.5). 58% of these were unidentified or uncharacterized in the blast search. GO annotation of the poorly annotated sequences suggests a high number of transcripts associated with the membrane cell component, calcium ion or protein binding function, and the protein glycosylation process. Unlike the DE analyses, these correlations do not account for tipping points in biological response, but are consistent across seasons. These genes likely are good targets for biological monitoring of acidification stress in pteropods, as they are expressed in laboratory exposures in multiple seasons in a consistent relationship to saturation state (Table S3). To develop any of these transcripts into an informative biomarker, we would, however, need to conduct a similar multi-season duration analysis as that which was conducted for shell quality (summarized in Figure 6), since gene expression has previously been demonstrated to be highly responsive to duration of exposure (Maas et al., 2018). These findings would then need to be validated with qPCR approaches to reach a similar level of technical and financial feasibility as shell condition analysis for implementation into routine monitoring. Additionally, the lack of full annotation of many of the most well correlated and consistently DE transcripts prevents us from having a functional understanding of their role in pteropod biology, pointing to areas of important further research.

Broadly, these findings demonstrate that seasonality plays a large role in the overall gene expression of *L. retroversa* and that there are interactive effects of CO_2_ and seasonality on the respiration rate of this species that appears to be linked to food availability or ontogeny. The response to CO_2_ exposure is, however, consistent among seasons for shell condition and, to a lesser degree, a subset of the transcripts. This suggests that there is specificity and repeatability in these metrics, making them viable tools for bioindicator implementation. The low cost, low-tech design of the shell condition workflow, as well as the fully developed intensity and duration characterization of the shell condition response, makes it an attractive choice for immediate implementation into biomonitoring and modeling efforts over the previously proposed pteropod bioindicator metrics (Bednaršek et al., 2017). Although prior work seems to indicate that the 1.5 aragonite saturation state threshold holds for multiple species of *Limacina* (Bednarsek et al., 2019), the quantitative relationships between saturation state and transparency should be explored in other species to determine the broader applicability of these equations.

## Acknowledgements

At sea sampling for this project was supported by Phil Alatalo, Leocadio Blanco Bercial, Sophie Chu, Taylor Crawford, Maja Edenius, Katherine Hoering, Ian Jones, Robert Levine, Mike Lowe, Camille Pagniello, Lenna Quackenbush, Andrea Schlunk, Ali Thabet, and Peter Wiebe. Captain K. Houtler and Mate I. Hanley made multiple seasons of sampling pleasant, safe, and effective aboard the R/V Tioga, and we note with deep sadness the passing of Capt. Houtler in 2021. We appreciate the efforts of Nancy Copley and Katherine Hoering in the lab for organizational and analytical assistance. Our gratitude for the patience of Leocadio Blanco Bercial for mathematical proofreading cannot be understated. We additionally acknowledge the importance of the conversations with Chris Melrose for refining our perspective on biological monitoring.

## Competing Interests

The authors have no competing interests.

## Funding

Funding for this project was provided by the National Science Foundation’s Ocean Acidification program (project OCE-1316040) to AEM, GLL, AMT and by NOAA Ocean Acidification Program Project # 21161 to AEM. Additional support for field sampling was provided by the WHOI Coastal Ocean Institute, the Pickman Foundation and the Tom Haas Fund at the New Hampshire Charitable Foundation to GLL and ZAW.

## Data Availability

All metabolic, shell condition and carbonate chemistry data are available at BCO-DMO (Project 2263). Gene expression data is available at Genbank (NCBI BioProject PRJNA260534).

## Author contributions

A.E.M lead the design of the 2014 cruises and experiments. She collected and analyzed the respiration and gene expression results, lead the interpretation of all datasets, and was responsible for the first draft of the paper. G.L.L contributed substantially to experimental design, and participated in all cruises and experiments. A.M.T. participated in cruises and experiments, and contributed substantially to all gene expression analyses. Z.A.W. contributed to experimental design, lead carbonate chemistry analysis and interpretation. A.J.B. participated in all cruises, ran the 2015 experiments, developed the shell transparency analysis, and made all shell transparency measurements. All authors contributed to data interpretation and to drafting and refining the manuscript.

---

**Figure S1:**
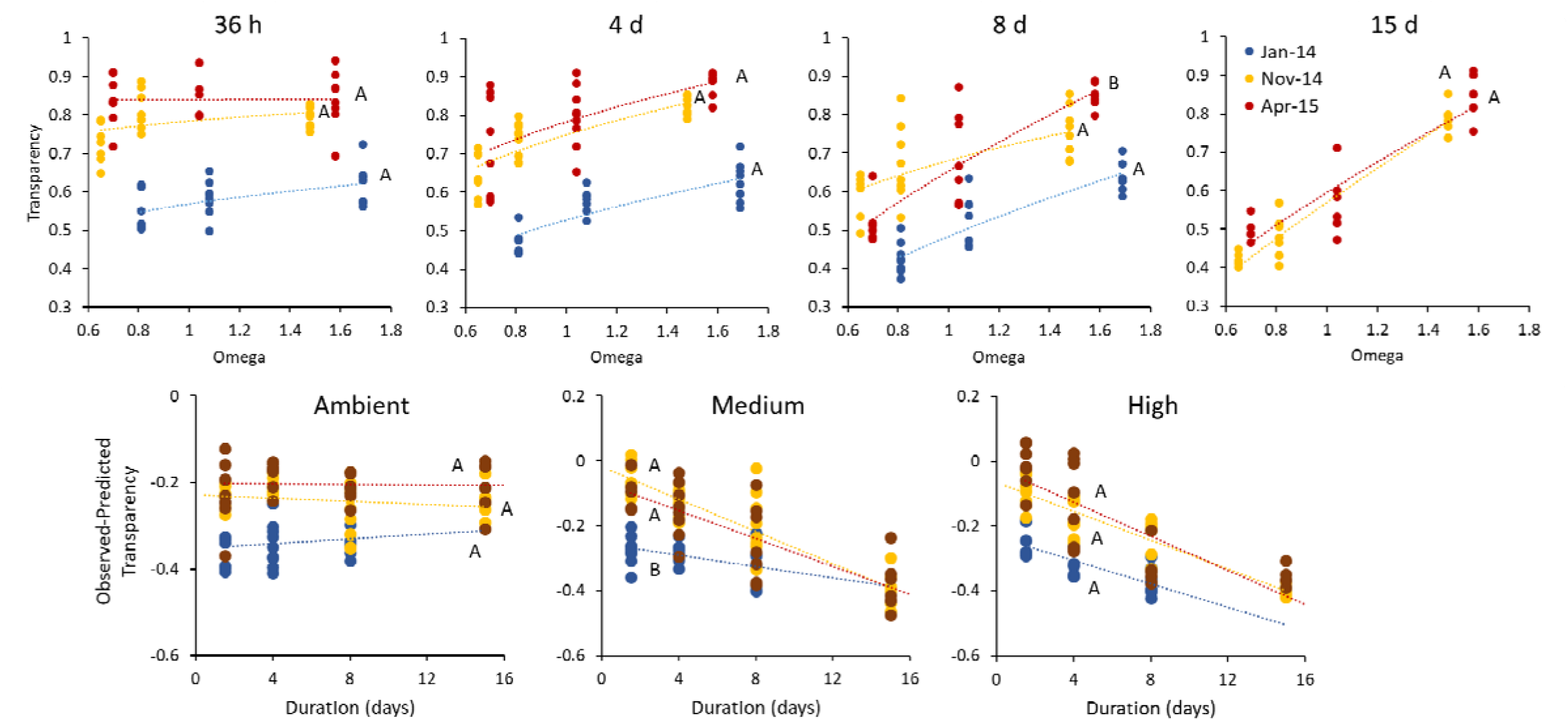
Detailed seasonal patterns of exposure duration effect on shell transparency. There was typically a consistent effect of the intensity of CO_2_ exposure (aragonite saturation state) on shell transparency during each day of the duration experiments (upper row), with the sampling period (color) having no significant effect on the slope of the relationship (power regression; letters denote statistically similar slopes based on Bonferroni post hoc analysis). During the 8 d duration analysis the slope of the relationship in April was statistically steeper. Analysis of the duration effect after correcting for saturation state (lower row) demonstrated no interactive effect between duration and sampling period (color), and suggests that there was no effect of duration on the ambient treatments. The slope of the relationship between duration and change in transparency was statistically similar in the medium and high treatments.

**Figure S2:**
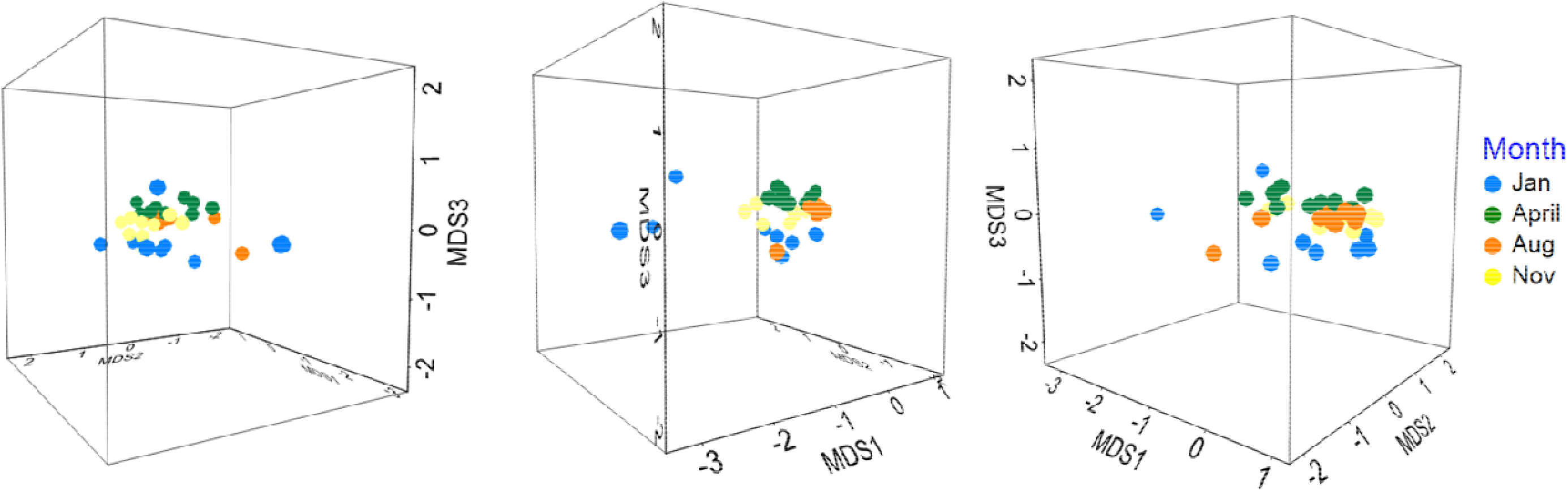
nMDS of Differential Gene Expression 3D. Three rotated perspectives of the third dimension of the DE nMDS plot (visualized in Fig. 4B) demonstrates the seasonal clumping of the data that is less obvious in the 2D version.

## Notes

### Competing Interest Statement

The authors have declared no competing interest.

